# Bright Probes, Blurred Metabolism: Navigating Fluorescent Protein Cross-Excitation in NADH FLIM

**DOI:** 10.64898/2026.05.27.727219

**Authors:** Janielle Cuala, Oscar Alberto de la Fuente, Lynne Cherchia, Yuanzhong Pan, Tvisha Singh, Chris Deng, Andrea Velazquez, Kate White, Senta K. Georgia, Steve Kay, Scott E. Fraser, Falk Schneider

## Abstract

Fluorescence lifetime imaging microscopy (FLIM) of endogenous NAD(P)H enables the label-free assessment of cellular metabolic state. Although metabolic imaging is increasingly combined with fluorescent protein (FP) reporters to enhance biological specificity, the potential cross-talk between the intrinsic and extrinsic labels remain ill-defined. Here, we systematically evaluate cross-talk from FPs in metabolic FLIM using phasor analysis of two-photon fluorescence microscopy. The results clearly show that many widely used fluorescent proteins are excited under the conditions used for NADH imaging; they emit blue-shifted, short-lifetime fluorescence that can interfere with imaging NADH metabolic signatures. This overlap persists across excitation wavelengths and FP classes, posing a significant challenge for multiplexed metabolic imaging. This cautionary tale argues against unvalidated multiplexing strategies in metabolic FLIM studies. Our study aims to identify acceptable imaging partners, offer a pipeline for assaying potential cross-talk, and provide practical guidance for experimental design.

**Significance:** Fluorescence lifetime imaging of NADH autofluorescence is a powerful, label-free approach to map cellular metabolism in living tissues. A growing number of studies combine NADH imaging with fluorescent protein (FP) reporters to simultaneously identify specific cell types or subcellular compartments. This study reveals that many FPs, spanning the visible spectrum, are unexpectedly excited under NADH conditions. Commonly used green, yellow, and red variants produce short-lifetime, blue-shifted fluorescence that directly overlaps with metabolic NADH signals. This cross-excitation can be falsely interpreted as a shift in cellular metabolic state, posing a significant risk for multiplexed metabolic imaging studies. Our studies establish a pipeline to assess and manage this risk. We identify StayGoldE138D and mNeonGreen as the most compatible FPs for co-imaging with NADH, and provide a practical framework to guide experimental design and control strategies for multiplexed metabolic FLIM.

## Introduction

Cellular metabolism rapidly adapts to changes in physiology, stress, and disease and is an indicator of cellular state. The ability to monitor metabolic states and their transitions therefore represents a powerful approach to study cellular function and diagnose disease^1,2^. Cancer cells, for instance, shift their metabolism towards glycolysis to sustain high ATP demand, known as the Warburg effect^3,4^. In the context of COVID-19 infection, cells such as monocytes and macrophages undergo metabolic changes during infection from the energy-efficient process of oxidative phosphorylation (OXPHOS) to glycolysis, which has been implicated in enhanced SARS-CoV-2 replication^5^. In diabetes, hyperglycemia reduces pancreatic beta cell mitochondrial metabolism, observed as an upregulation of glycolysis-related genes and a reduction of OXPHOS^6^. Together, these examples highlight metabolism as both a readout and a driver of disease progression.

Robustly measuring changes to cellular metabolism is of critical importance to understanding cellular physiology. Biochemical approaches can be used to monitor biomarkers and extracellular acidification^7^, however, such methods rely on homogeneous samples or dissociated tissues. Microscopy methods, instead, enable non-invasive observation of changes to metabolism in intact and living samples. To this end, biosensors for visualizing specific metabolites have been developed^8^. For instance, PercevalHR can be used to look at changes in ATP/ADP ratio^9^, FapR-NLuc to observe Malonyl-CoA in liver fatty acid metabolism^10,11^, TINGL to monitor glucose levels^12^, and Peredox to report on NADH/NAD+ ratio^13^. While increasingly powerful, such biosensors’ potential can be limited by experimental variables such as expression level, detection efficiencies, varying dissociation constants, or changes within the local physico-chemical environment.

Metabolic imaging exploits the autofluorescence of endogenous metabolic compounds and thus presents a label-free remedy. Specifically, NAD(P)H and FADH are intermediate substrates in metabolism that can be excited using ultraviolet light or two-photon excitation (TPE)^14,15^. Their concentration and interaction with enzymes of metabolic pathways (**Fig. 1a**) are important factors in assessing the metabolic state of cells and tissues^16^. In the following, NADH will be used to collectively refer to both NADH and NADPH. Importantly, the interaction of NADH with enzymes causes only minor changes to its emission spectrum but results in measurable changes to its fluorescence lifetime, which is the average time a fluorophore spends in the excited state. This is used in fluorescence lifetime imaging microscopy (FLIM) as an additional means to generate contrast in the image. The fluorescence lifetime of free NADH is ∼0.4 ns and increases when bound to different enzymes to 1-5 ns (see illustration in **Fig. 1b**). The longer the lifetime, the more NADH is bound to proteins, specifically to enzymes of the OXPHOS pathway^15,17^. In this way, changes to NADH lifetime can be used as a biophysical readout to report on shifts in cellular metabolism from glycolysis to OXPHOS or vice versa^18^.

**Figure 1:**
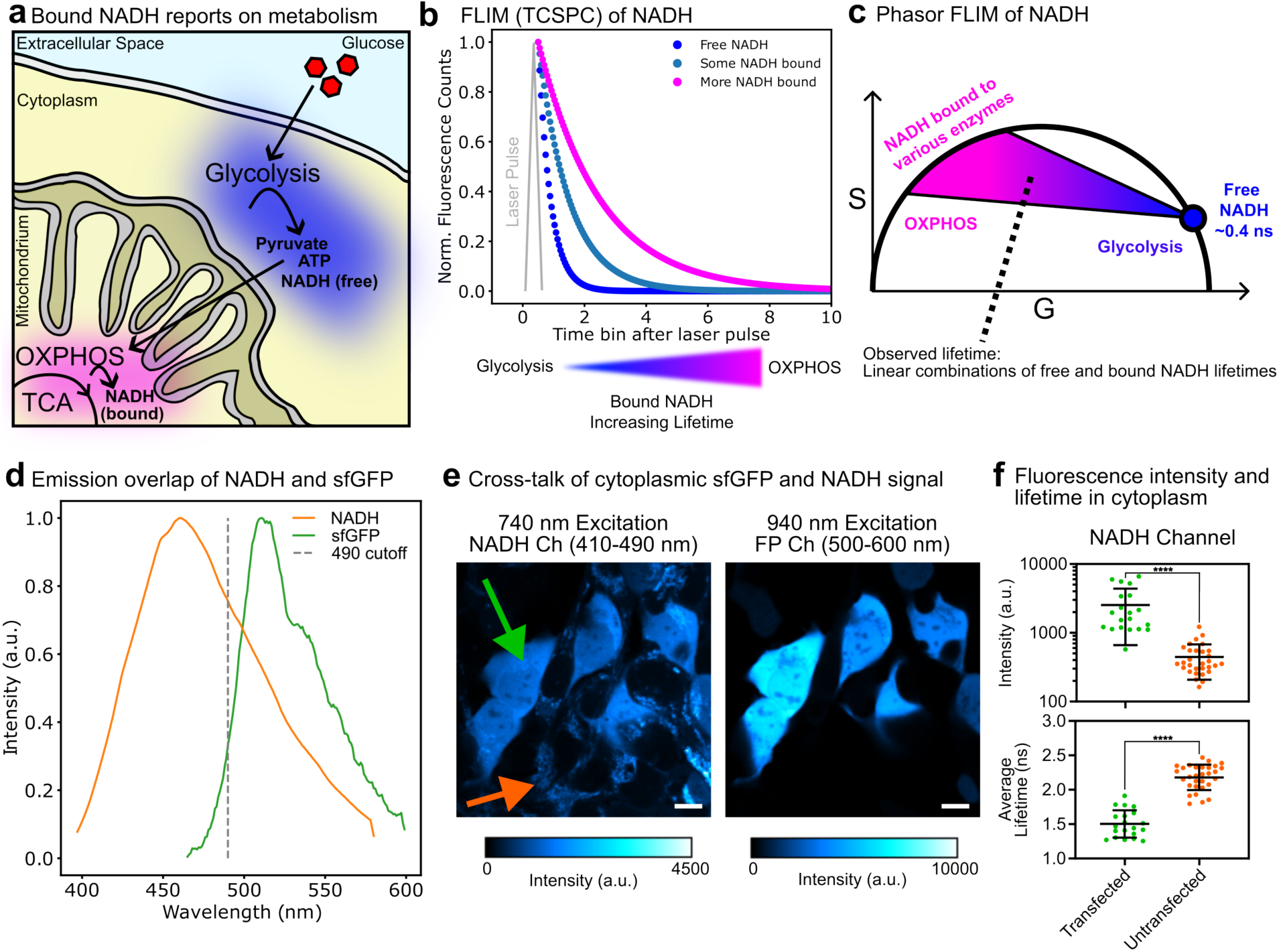
Metabolic imaging with FLIM. **a** Cartoon illustrating subcellular localization of glycolysis and oxidative phosphorylation (OXPHOS), as well as resulting bound or free NADH. **b** Exponential decays illustrating time correlated single photon counting decays of free NADH (assuming a lifetime of 0.4 ns) and bound NADH (longer lifetimes). **c** Cartoon of a phasor plot for metabolic imaging. Monoexponential lifetimes fall on the unity circle. Free NADH has a monoexponential lifetime of 0.4 ns. NADH bound to various enzymes (mixed with free NADH) can be found on the metabolic trajectory indicated by blue to magenta shading. Shorter lifetimes towards free NADH (blue) indicate glycolysis, whereas longer lifetimes (magenta) indicate OXPHOS as the prevalent metabolic state in a measurement. **d** Normalized fluorescence emission spectra of NADH and sfGFP-H2B expressed in HEK cells. **e** Laser scanning two-photon excitation microscopy images of HEK cells expressing cytoplasmic sfGFP. With 740 nm excitation and NADH detection 410-490 nm (left) and FP detection 500-600 nm (right), mitochondria and cytoplasmic sfGFP can be observed. Arrows indicate transfected and untransfected cells (green and orange, respectively). With 940 nm excitation and FP detection, sfGFP can be observed. Scale bars are 10 µm. **f** Fluorescence intensity and average fluorescence lifetime of the cytoplasm area segmented from the transfected and untransfected cells. Every dot represents an area segmented from a single cell. Horizontal bars indicate mean values and vertical lines indicate standard deviations (p ≤ 0.0001 = ****, from two-tailed t-test).

Fluorescence lifetime data can be acquired using one-photon excitation (OPE) or TPE. Due to the superior signal-to-noise ratio (SNR), higher penetration depth, and reduced phototoxicity compared to UV exposure, TPE is commonly used to study free and bound NADH levels^14,18^. It can be combined with FLIM acquisitions in the frequency domain using excitation phase modulation or using pulsed excitation with time-correlated single photon counting (TCSPC), leading to fluorescence decay curves per pixel (**Fig. 1b**). For the latter, decay data are typically fitted by exponential models wherein the decay time represents the lifetime. In the case of metabolic imaging data, fitting can be a difficult task as mixtures of species (free NADH and NADH bound to various enzymes) are present and vary spatially. The phasor approach to fluorescence lifetime presents a remedy as it converts the complex modulation or decay data to a projection on a 2D plane (**Fig. 1c**)^19^. Every pixel in the FLIM dataset has an intensity value, an average lifetime value, and two phasor coefficients, usually labelled G and S. The phasor approach does not require fitting and thus does not make assumptions about the number of components present. It does indicate if the decay is mono- or multi-exponential as single component data map onto the phasor circumference, whereas multicomponent data fall within the circle. Furthermore, it helps deal with low SNR data and explore the number of different lifetime components in the sample^20–22^. Phasor analysis is becoming the go-to tool for metabolic imaging.

Label-free FLIM can be combined with fluorescent labels to increase the information content and provide cellular context. For example, labeling a certain population of cells (e.g., cancerous, infected, diabetic) with an additional fluorophore can provide mechanistic insights. The fluorophores of choice are typically fluorescent proteins (FPs), which can be conveniently expressed in the cells of interest. Metabolic imaging of NADH has been reported in combination with imaging of, for example, GFP^18^, EGFP^23,24^, or mCherry^25^. Fluorescence emission from the FPs and NADH was reported to be separated using stringent emission filters. However, concerns regarding cross-excitation of NADH and FPs, signal spill-overs, and blue-shifted FP emission have been raised^26^. The photophysics of FPs are complicated, depending on the exact sequence of the FP and the cellular environment, and might vary between OPE and TPE^27–30^. Thus, different versions of EGFP could have completely different TPE properties. Due to the dim nature of NADH fluorescence (low quantum yield) as compared to FPs, any cross-talk would be particularly difficult to account for.

In this work, we systematically address the issue of metabolic imaging of NADH in the presence of another fluorescently labelled species. We employ the phasor approach to separate NADH from FP signal and reveal blue-shifted fluorescence with short lifetime for multiple fluorescent proteins, which can potentially pollute the metabolic imaging of NADH. We demonstrate that all tested FPs show overlap with the metabolic imaging space mapped out by drug and glucose treatments. Overall, we aim to provide comprehensive guidelines for performing metabolic imaging in context.

## Materials and Methods

### Reagents

**Table 1:**
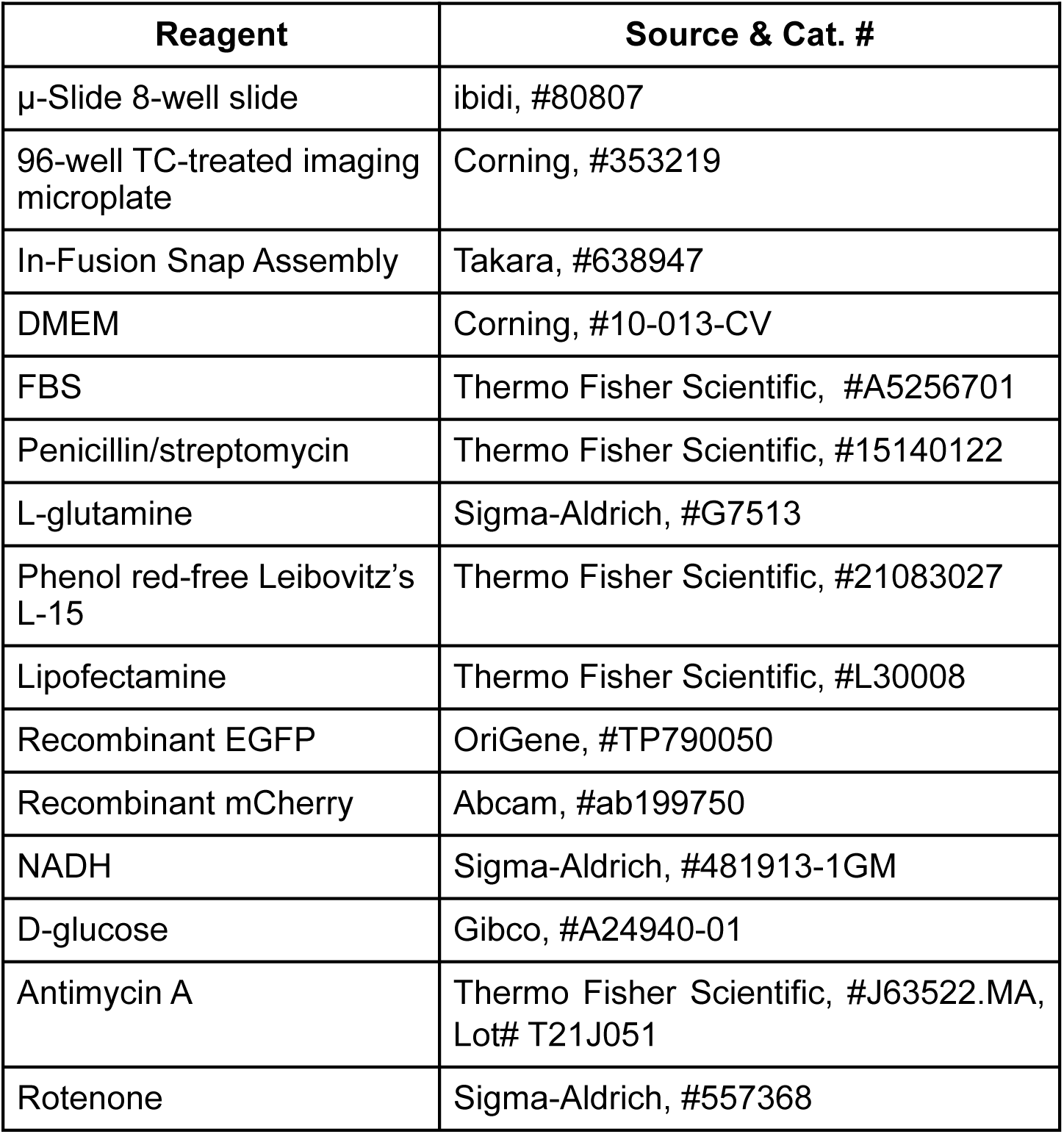
Reagents used in this study.

| Reagent | Source & Cat. # |
| --- | --- |
| μ-Slide 8-well slide | ibidi, #80807 |
| 96-well TC-treated imaging microplate | Corning, #353219 |
| In-Fusion Snap Assembly | Takara, #638947 |
| DMEM | Corning, #10-013-CV |
| FBS | Thermo Fisher Scientific, #A5256701 |
| Penicillin/streptomycin | Thermo Fisher Scientific, #15140122 |
| L-glutamine | Sigma-Aldrich, #G7513 |
| Phenol red-free Leibovitz's L-15 | Thermo Fisher Scientific, #21083027 |
| Lipofectamine | Thermo Fisher Scientific, #L30008 |
| Recombinant EGFP | OriGene, #TP790050 |
| Recombinant mCherry | Abcam, #ab199750 |
| NADH | Sigma-Aldrich, #481913-1GM |
| D-glucose | Gibco, #A24940-01 |
| Antimycin A | Thermo Fisher Scientific, #J63522.MA, Lot# T21J051 |
| Rotenone | Sigma-Aldrich, #557368 |

### Plasmid design

All plasmids used in this study are summarized in Table 2. For the TPE cross-talk assay, all FPs have been fused to human H2B, creating a library of constructs to test. The FP amino acid sequences have been checked against the respective entry on FPbase.org^31^. The H2B sequence has been PCR amplified from a pCS2+-mCherry-H2B plasmid (kind gift from Le Trinh, USC). PCRs were performed using LA Taq (#RR002A) according to the manufacturer’s protocol. A short linker sequence has been introduced to separate FP and H2B: GSGPVATGSG. Cloning has been performed using Takara In-Fusion Snap Assembly (#638947) according to the manufacturer’s protocol. We used a pCS2+ plasmid (kind gift from Le Trinh, USC) containing a CMV promoter for expression in mammalian cells as a backbone. sfGFP-N1 was a gift from Michael Davidson and Geoffrey Waldo (Addgene plasmid # 54737; http://n2t.net/ addgene:54737; RRID:Addgene_54737). FPs were obtained as gene blocks with flanking regions for In-Fusion (synthesized by IDT) or amplified by PCR. The enhancer vhh sequence was taken from prior studies^32,33^. A T2A sequence (EGRGSLLTCGDVEENPGP) has been used for equimolar expression of vhh and FP-H2B^34^.

**Table 2:**
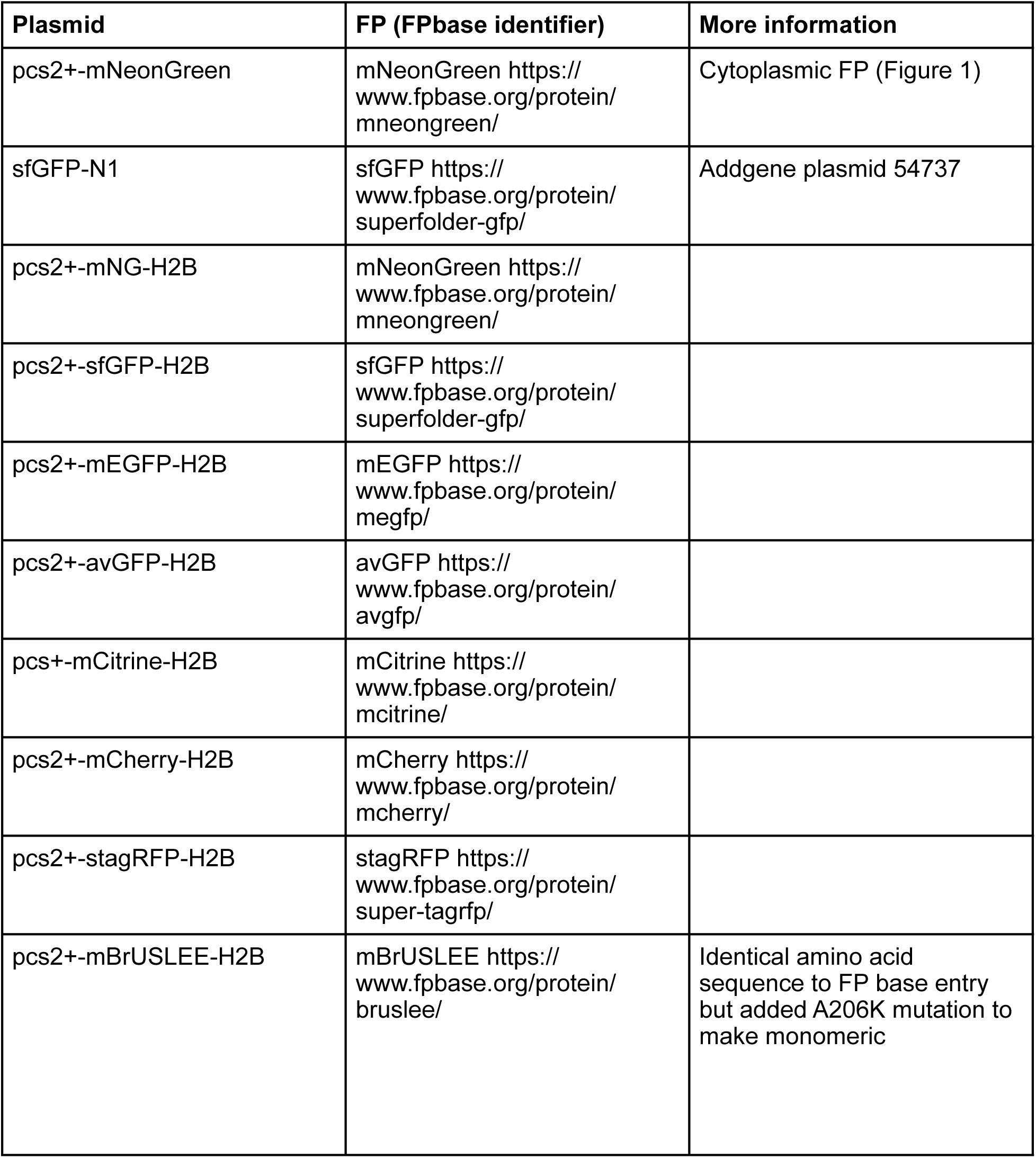
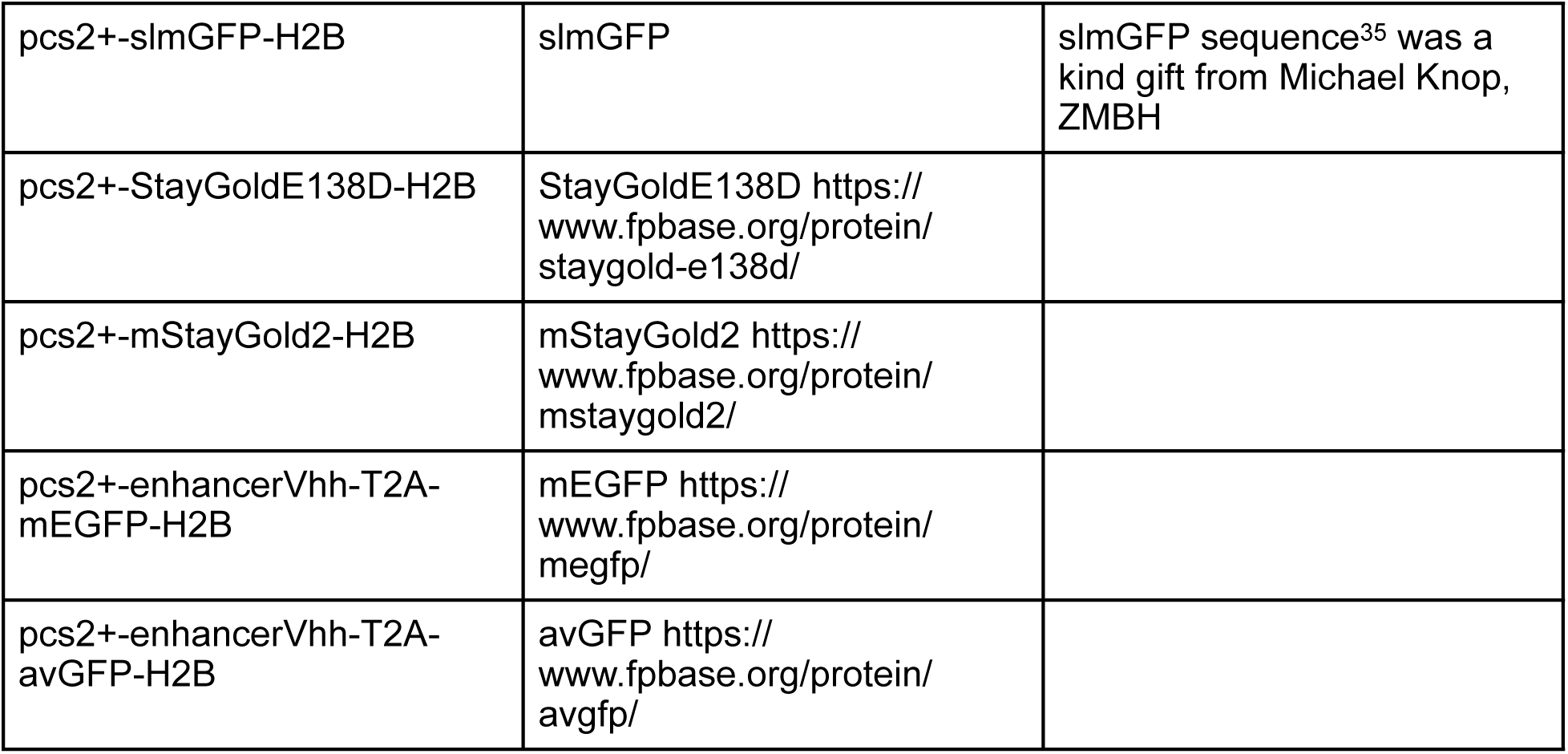
Plasmids used in this study.

### Cell culture and transfection

U2OS cells were obtained from the American Type Culture Collection and cultured in DMEM (Corning, #10-013-CV) supplemented with 10% fetal bovine serum (Thermo Fisher Scientific, #A5256701), 1% penicillin/streptomycin (Thermo Fisher Scientific, #15140122), and 1% L-glutamine (Sigma-Aldrich, #G7513). Cells were grown to approximately 80% confluency before being seeded onto a µ-Slide 8 well (ibidi, #80807). 24 h after seeding, cells were transfected using Lipofectamine 3000 and P3000 reagents (Thermo Fisher Scientific, #L30008) according to the manufacturer’s protocol. 24-48 h after transfection, cell culture medium was replaced with prewarmed, phenol red-free Leibovitz’s L-15 medium (Thermo Fisher Scientific, #21083027) prior to image acquisition.

### Plate Reader: Spectra

U2OS cells were seeded onto a black 96-well plate (Corning, #353219) and transfected as described above. 24 h after transfection, cell culture medium was replaced with prewarmed L-15, followed by a spectra scan on a CLARIOstar Plus microplate reader using CLARIOstar MARS software. Spectra were exported to Excel reports, averaged, and smoothed over 3-6 technical replicates using custom Python code (https://github.com/janiellecuala/Multiplexing-NADH-and-Fluorescent-Proteins-in-Metabolic-Imaging).

### Two-Photon Excitation Microscopy

Fluorescence Lifetime Imaging Microscopy (FLIM) was performed on an SP8 DIVE FALCON spectral multi-photon FLIM microscope (Leica Microsystems, Germany) using an HC PL AP 63x/1.2NA motCorr water immersion objective (Leica, #11506361). NAD(P)H was excited with a Spectra-Physics Insight 3X ultrafast IR laser, 0.8 mW average power, and 20-40 frame accumulations per optical section. Excitation was performed as indicated: 740 nm was used for NADH excitation, 940 nm was used for fluorophore excitation, except for stagRFP and recombinant mCherry, which were excited at 1050 nm. Most images were collected at 512 x 512 pixels and 2.0-2.5 zoom, resulting in a final pixel size of 144 nm/px. Spectral detection was performed using SMD-HyD detectors in non-descanned (NDD) configuration, with detection windows as indicated: 410-490 nm for NADH, 500-600 nm for green/yellow fluorescent protein detection, and 600-700 nm for red fluorescent protein detection. Where indicated, lambda scans were performed by varying the excitation or emission wavelength by 10 nm increments.

In vitro images were collected with 20 frame repetitions at 128 × 128 pixels, using a zoom factor of 5, which resulted in a final pixel size of 290 nm/px.

### Metabolic trajectory and in vitro measurements

Glucose-stimulated experiments were performed with solutions made from Krebs-Ringer Buffer solution to make 2.8 mM low glucose (LG), 16.7 mM high glucose (HG) (Gibco, #A24940-01), and HG with 1 µM rotenone (Sigma-Aldrich, #557368) and with 1 µg/mL antimycin-A (HG+Drug) (Thermo Fisher Scientific, #J63522.MA). All samples were starved in LG solution for 1 h; the LG condition was then imaged. For the HG condition, the LG solution was replaced with HG solution for 30 minutes and then imaged. Finally, for the HG+drug condition, the HG solution was then replaced with the HG+drug solution for 30 minutes and then imaged.

For in vitro measurements, recombinant proteins and NADH were imaged in L-15 on µ-Slide 8-well coverslips. EGFP (OriGene, #TP790050) was imaged at a concentration of 150 nM, mCherry (Abcam, #ab199750) was imaged at 250 nM, and NADH (Sigma-Aldrich, #481913-1GM) was imaged at 200 µM.

### Data processing and analysis

FLIM images were preprocessed in the Leica LAS X software using the phasor FLIM module. Pixels are binned 3 by 3, and a threshold of 10 photons was applied (i.e., all pixels with less than 10 photons are disregarded). Phasor coordinates were filtered using the built-in wavelet filter. Fast FLIM images were exported along with intensity and G/S images. Thus, for each detector, a total of 4 images were exported: from channels 0 to 3, representing intensity, lifetime, G, and S coordinates, respectively. Any pseudocolored lifetime images were directly exported from LASX. For pseudocolored metabolic images, the LASX metabolic arc module was applied to the phasors, with free NADH set at 0.4 ns.

Tiff files were processed using Fiji/Image J^36^. ROIs for transfected nuclei were drawn based on the intensity image of the fluorophore; untransfected nuclei and cytoplasm ROIs were selected based on the NADH intensity image. Separate CSV files were exported with the mean values for intensity, lifetimes, and G and S coordinates. A macro for automated processing in FIJI can be found on our GitHub.

For excitation and emission scan plots, mean intensity values from the exported CSV file for the ROIs from 680 nm to 780 nm, normalized to global maximum per FP, were plotted through the Python code found on our GitHub. For metabolic trajectory boundary analysis, mean G and S coordinates were plotted with Prism 10. For each experiment, at least 3 biological replicates (i.e., different imaging sessions with various sets of cells) were done, and at least 5 technical replicates (i.e., number of transfected, untransfected, and cytoplasm ROIs) for each fluorescent protein or condition.

## Results

We set out to multiplex NADH and fluorescent proteins on a TPE setup. We began by assessing green FPs, as the OPE emission spectra of sfGFP, avGFP, mEGFP, and mNeonGreen could be reasonably separated from NADH (**Supp. Fig. S1**), allowing for the use of further red-shifted probes to label additional biological structures. The combination of sfGFP and NADH, in particular, would represent a powerful pair for metabolic imaging in the context of diabetes research due to the existence of the c-peptide-sfGFP (CpepSfGFP) mouse line^37^. The OPE emission spectra of NADH and sfGFP show only minor overlap (**Fig. 1d**). Cross-talk from NADH to the FP detection channel is less of a concern due to the difference in brightness of the fluorophores, but cross-talk from sfGFP to the NADH channel would pose an issue. However, stringent filters, different excitation wavelengths, and dedicated tracks may allow simultaneous imaging. When employing TPE at 940 nm on cells transfected with sfGFP, we can clearly identify the FP fluorescence (in FP emission channel, 500-600 nm). When using 740 nm for excitation, we do not expect excitation of sfGFP. However, cross-talk becomes apparent in fluorescence images as cytoplasmic and nucleoplasmic haze, while untransfected cells only show mitochondrial signal from NADH autofluorescence (**Fig. 1e**). At 740 nm excitation, the raw intensity is about five times higher in transfected versus untransfected cells, and a decrease in average fluorescence lifetime, from 2.2 ns to 1.5 ns, is observed (**Fig. 1f**), Indicative of a shift in metabolic state towards glycolysis. As sfGFP shows fluorescence emission overlap with NADH emission, we assessed cytoplasmic mNeonGreen in the same way and observed distinct cytoplasmic haze in transfected cells, albeit less pronounced than for sfGFP (**Supp. Fig. S2**). To assess green/yellow FP emission in the absence of the added complexity of cells, we recorded emission spectra of recombinant EGFP excited at 740 nm and 940 nm (TPE) and observed extension of the emission peak to shorter wavelengths under 740 nm TPE (**Supp. Fig. S3**). Taken together, these experiments lead us to believe that some green fluorescent proteins can be excited using 740 nm and show unusual fluorescence properties such as blue-shifted emission and altered lifetime. This may cause cross-talk with the NADH signal, potentially biasing metabolic imaging.

These issues are challenging to assess in live cells expressing cytoplasmic FPs due to spatial overlap with NADH fluorescence (cytoplasmic and mitochondrial). To investigate cross-talk under cellular/physiological conditions, we designed an assay with nuclear-localized fluorescent protein (conjugated to H2B) to better separate NADH and FP fluorescence, as only minimal signal from free NADH is expected in the nucleus^38–40^ (**Fig. 2a**). We recorded fluorescence at 740 nm and 940 nm TPE in the NADH channel (410-490 nm) and FP channel (500-600 nm). The resulting four-channel images and respective FLIM data are displayed and analyzed using phasor plots and inform on fluorescence properties and cross-talk (**Fig. 2a**). Ideally, the FP should show close to monoexponential lifetime and map to the phasor circle when excited with 940 nm (emission is expected roughly between 500-600 nm based on OPE spectra, **Supp Fig. 1**). Excitation with 740 nm should cause either no fluorescence or only weak fluorescence in the FP channel. NADH autofluorescence should not be observed in the FP channel, and FP fluorescence should not show in the NADH channel. Thus, the four phasor plots can be used to assess potential cross-excitation and cross-emission of FP and NADH quickly, and should allow for observation of blue-shifted fluorescence, if present.

**Figure 2:**
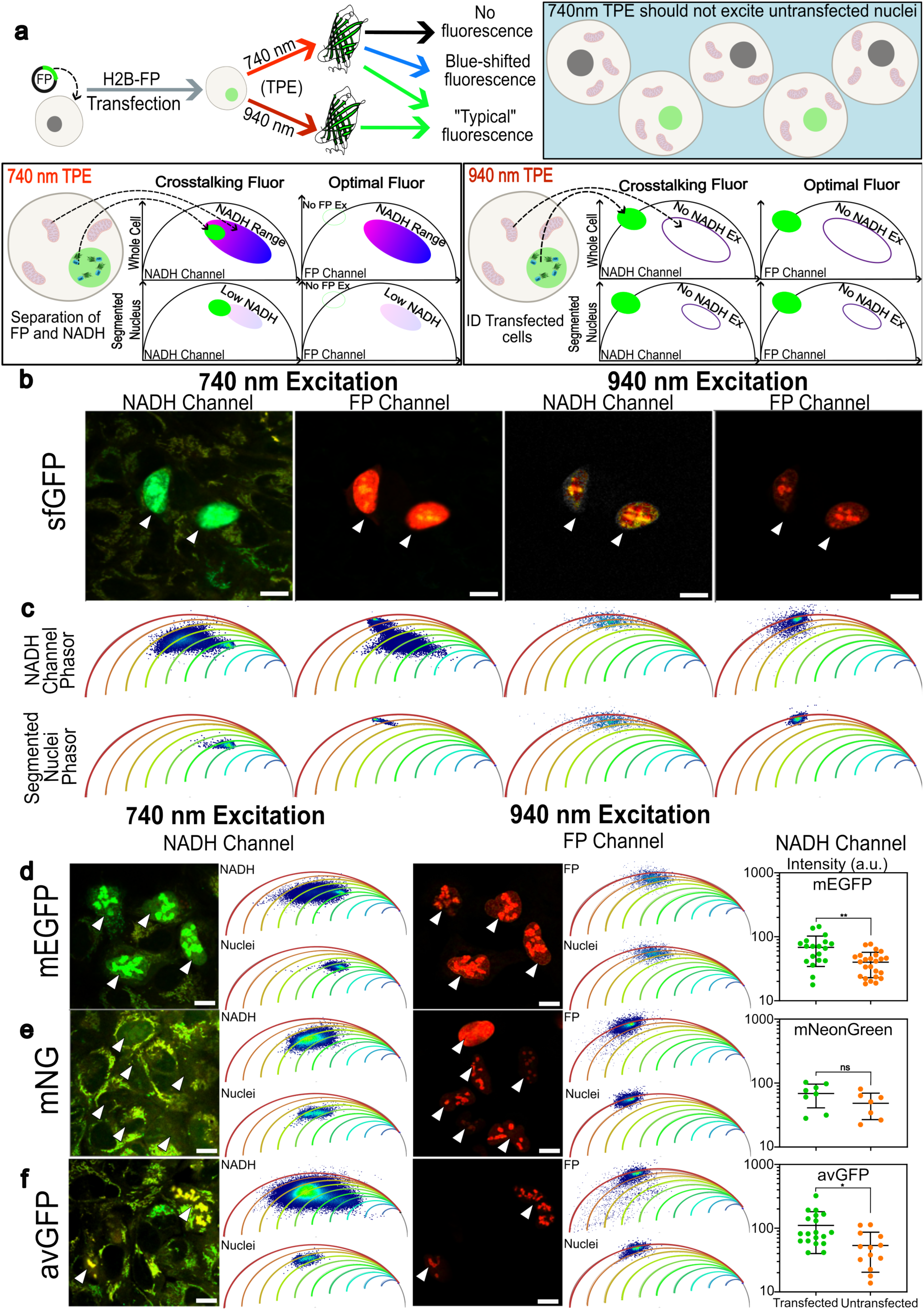
TPE cross-excitation assay using nuclear-localized FPs. **a** Cartoon illustrating experimental setup of TPE cross-excitation assay. Briefly, U2OS cells are transfected with the respective fluorescent protein (FP). Using TPE excitation at 740nm, the FP can be determined to be crosstalking or optimal based on phasor signal separation of FP and NADH signals. With TPE 940nm excitation, the identification of transfected nuclei can be confirmed. Ideally, nuclear localization of fluorophore expression minimises spatial crosstalk between FP and NADH signal. **b,c** TPE cross-excitation assay for sfGFP. **b** Laser scanning TPE microscopy images of U2OS cells expressing nuclear sfGFP with 740 nm excitation or 940 nm excitation. Detection in the NADH channel range 410-490 nm and FP range 500-600 nm. Arrows indicate the transfected cells with nuclear sfGFP. With 940 nm excitation and FP detection, sfGFP can be observed. Color is based on the phasor plot metabolic trajectories below. Scale bars are 10 µm. **c** Phasor plots associated with each image above are shown with metabolic arcs represented with a rainbow LUT to represent percent bound NADH, where blue is 0% and red is 100%. The top row represents the NADH channel that includes all pixels from the whole image. Bottom row is the phasor signal from the segmented nuclei of the transfected cells, shown with the white arrows from **b**. **d,e,f** TPE cross-excitation assay. Laser scanning TPE microscopy images of U2OS cells expressing nuclear mEGFP, **e** mNG, **f** avGFP. With 740 nm excitation, NADH channel detection is within the 410-490nm range. With 940nm excitation, FP channel detection within the 500-600nm range. Arrows indicate the transfected cells with respective nuclear FP. Similar to (c), phasor plots correspond to the images to its left, where the top phasor represents the NADH channel that includes all pixels from the whole image. Bottom row is the phasor signal from the segmented nuclei of the transfected cells, shown with the white arrows. The same rainbow LUT was used, and all scale bars are 10 µm. Intensity values of transfected and untransfected nuclei are plotted on the right of the respective FP. Every dot represents a segmented nucleus. Horizontal bars indicate mean values and vertical lines indicate standard deviations (p ≤ 0.05 = *, p ≤ 0.001 = **, from two-tailed t-test).

Performing the cross-excitation assay on sfGFP-H2B transfected cells reveals clear nuclear fluorescence from 740 nm and 940 nm TPE in all four channels (**Fig. 2b**). A low level of background fluorescence from the nuclei in the NADH channel is expected from the emission spectra (**Fig. 1d**). Fluorescence intensity images are color-coded using the phasors (isolines) to indicate metabolic ratios of the signal, referred to as a Phasor-FLIM image (**Fig 2b**). Phasor plot analysis of the whole images and segmented nuclei is shown in **Fig. 2c**. Transfected cells can easily be identified using the 940 nm excitation and the FP channel (500-600 nm emission). Here, we observe the typical fluorescence of sfGFP, with all pixels mapping close to the phasor unity circle. When excited with 740 nm, fluorescence from the transfected nuclei in the NADH channel (410-490 nm) is distinguishable from untransfected cells. This is indicative of cross-talk and confirms our previous observations that sfGFP excited at 740 nm shows blue-shifted fluorescence similar to the recombinant EGFP in solution (**Supp. Fig. S3)**. This analysis further demonstrates that the phasor coordinates from sfGFP in the NADH channel under 740 nm excitation are located inside the unity circle and thus show unusual properties (short, multi-component lifetime). This is distinctively different from emission cross-talk when the FP is excited with 940 nm and observed in the NADH channel. In that case, the nuclear pixels map close to the unity circle on the phasor, indicating monoexponential decay, which could easily be unmixed from the NADH fluorescence (**Fig. 2c**). Thus, fluorescence from sfGFP when multiplexed with NADH imaging could be mistaken for a metabolic shift towards glycolysis.

The fluorescence cross-talk from sfGFP might be a particular property of this FP. To investigate if this is a more general property of green fluorescent proteins, we performed the same experiments with avGFP (wildtype GFP from *acquoria victoria*), mEGFP, and mNeonGreen (mNG). Phasor-FLIM images and respective phasor plots are shown in **Fig. 2d-f**. When excited with 940 nm, each FP fluoresces with the expected typical properties (lifetime of 2-3 ns and mapping onto the phasor unity circle). However, when excited with 740 nm, all three green, nuclear-localized FPs fluoresce with a shortened lifetime, and their phasor coordinates overlay with the NADH autofluorescence phasor cloud. This is shown by segmenting nuclei of transfected cells and visualizing the respective phasor plots of these pixels (**Fig. 2d-f**). Furthermore, we note an increased background intensity of nuclei from transfected vs untransfected cells. Quantification of 740nm excitation-induced nuclear fluorescent intensity over the 410nm to 490nm emission range shows a statistically significant increase in signal for nuclei expressing avGPF and mEGFP (p-values 0.0103 and 0.0010, respectively), indicating significant FP excitation across the NADH detection range. However, while mNG-expressing nuclei show a slight increase in fluorescent signal over untransfected nuclei, the effect appears to not be statistically significant (p-value 0.1387). These results show that while the tested green fluorescent proteins all share some emission in the NADH range upon 740nm TPE, the degree of interference differs across FPs, therefore indicating that unique green FPs could prove to be better candidates for combined metabolic imaging due to decreased/negligible excitation under NADH imaging conditions. Each of these FPs might impede multiplexed metabolic imaging. Notably, avGFP shows less of a shift of the phasor cloud when comparing 740 and 940 nm excitation, which could be a result of the blue-shifted fluorescence excitation peak at about 400 nm compared to the other green FPs with excitation peaks around 490 nm (see OPE spectra in **Supp Fig. S1**). As fluorescence lifetime is a function of the local environment of the fluorophore, we confirmed our observations of blue-shifted short lifetime fluorescence (mapping towards the center of the phasor plot) in the cellular assay using recombinant, purified EGFP in solution (**Supp. Fig. 4a**).

For excitation and detection of NADH, various wavelengths have been used in the literature (**Table 3**). To investigate the effect of the TPE wavelengths on fluorescence cross-talk from the FPs, we performed an excitation scan varying excitation wavelengths from 680 nm to 780 nm and observed fluorescence in the NADH channel (**Fig. 3**). We segmented transfected nuclei, untransfected nuclei, and cytoplasm/mitochondria to compare fluorescence across TPE wavelengths. Transfected nuclei appear with higher mean fluorescence as compared to untransfected controls within the same image. Fluorescence from untransfected nuclei originates from low levels of free NADH in the nuclei. avGFP, mEGFP, and sf GFP show signal levels clearly above background (**Fig. 3a-c**). Nuclear-localized fluorescence from mNG is slightly lower and closer to the background (**Fig. 3d**) consistent with the cytoplasmic mNG (**Supp. Fig. 2**). Given the NADH and FP two-photon excitation profiles in the 680nm to 780nm range, it is difficult to disentangle their respective contributions to total intensity and apparent lifetime. In our hands, 740nm TPE offers a compromise.

**Figure 3:**
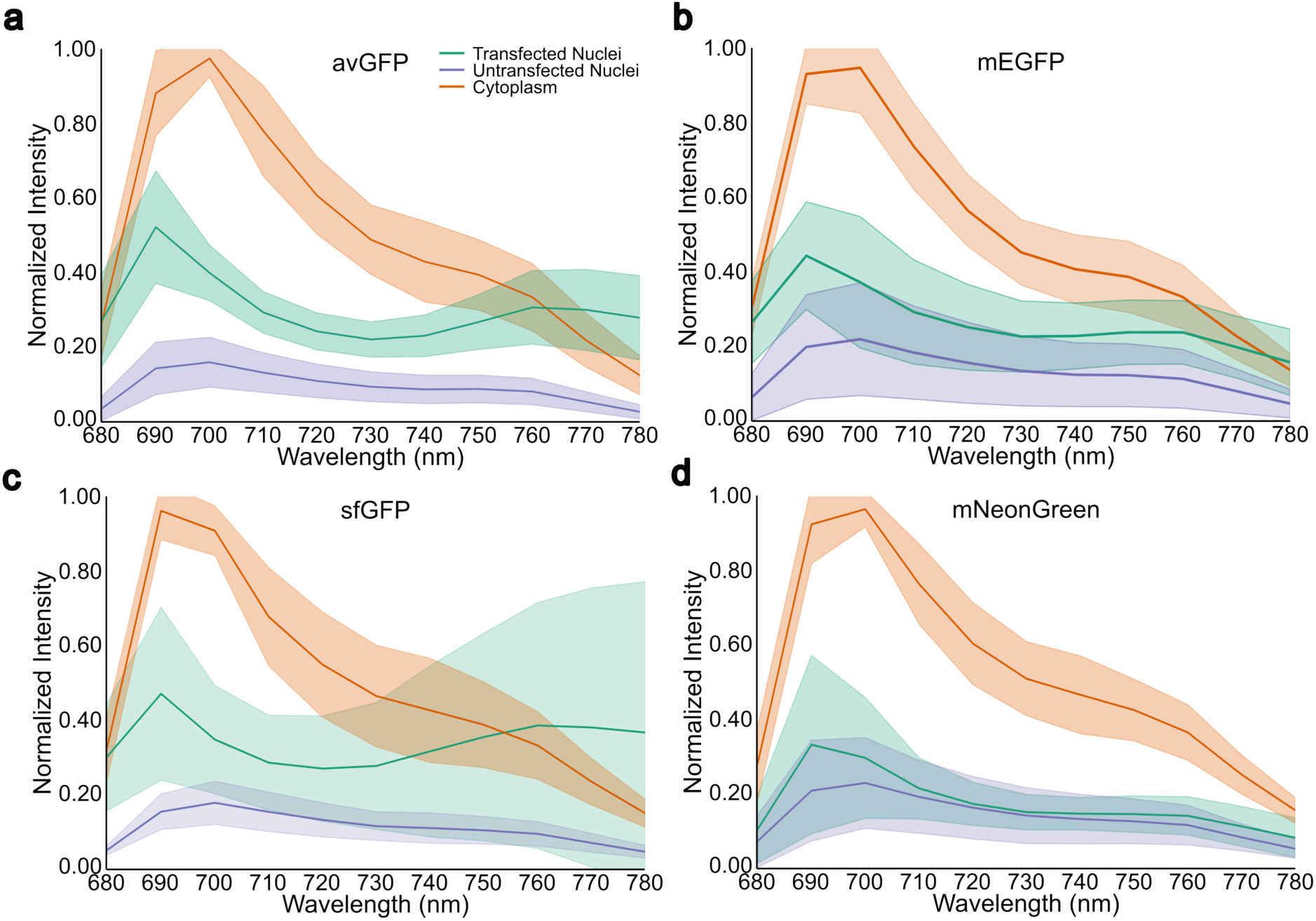
Two-photon excitation scan. Fluorescence intensity measurements from cytoplasm (orange line), untransfected control nuclei (blue line), and transfected nuclei (green line) of U2OS cells across excitation wavelengths from 680-780 nm with 10nm intervals in between. All measurements were normalized to excitation laser power (see Materials and Methods). Data points represent mean fluorescence intensities, and shaded regions indicate standard error of the mean. **a** avGFP, **b** eGFP, **c** sfGFP, **d** mNG. Each data point represents the average of measurements from at least 5 nuclei per condition across multiple biological replicates.

**Table 3:**
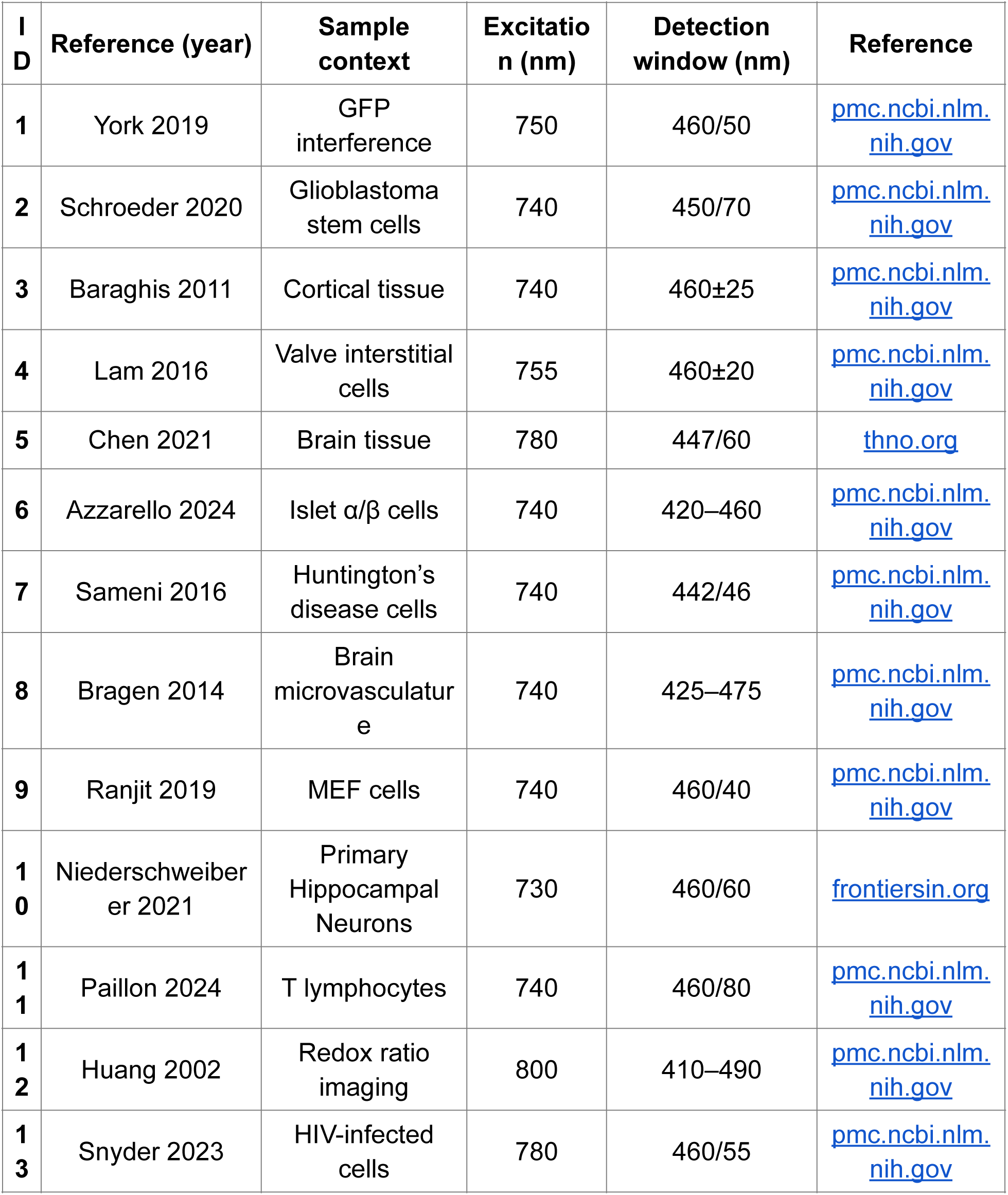
Example excitation and emission parameters for metabolic imaging of NADH reported in the literature.

Due to the cross-talk between 740 nm excited-green FPs and NADH, we sought to use bathochromic-shifted FPs, expecting to see less excitation cross-talk. We chose three FPs with increasing excitation maxima, mCitrine, stagRFP, and mCherry (516 nm, 555 nm, and 587 nm), whose OPE excitation and fluorescence spectra are well separated from the NADH channel (**Supp. Fig. 5**). We utilized our TPE assay with H2B-FP fusions to experimentally test for cross-talk with metabolic imaging (**Figure 4**). For all three red-shifted FPs, we observed excitation at 740 nm and fluorescence in the NADH channel. The segmented nuclei of transfected cells are clearly discernible on the Phasor-FLIM images and from the NADH cloud on the phasor plot, but might be mistaken for metabolic shifts towards glycolysis. As mentioned above, we performed a TPE scan from 680 nm to 780 nm and observed above-background fluorescence in the nuclei for all three FPs (**Figure 4**). mCherry shows the least excitation and fluorescence in the 680-780 nm range as compared to untransfected background (**Figure 4c**). We verified the TPE of mCherry at 740 nm and the blue-shifted fluorescence in vitro with purified recombinant protein (**Supp. Fig. 4B** and **Supp. Fig. 6**).

**Figure 4:**
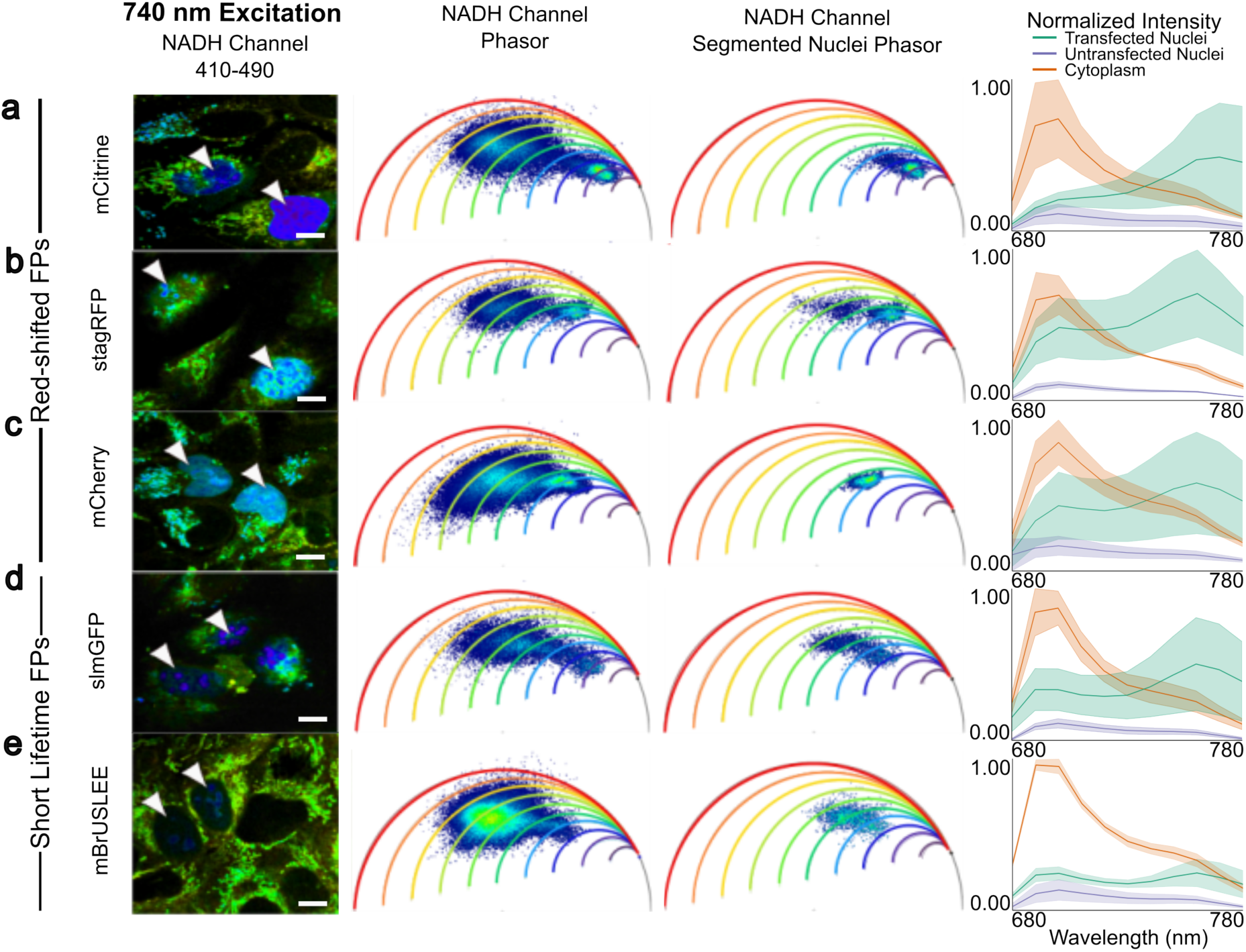
TPE cross-talk assay on bathochromic and short lifetime FPs. Laser scanning TPE microscopy images of U2OS cells expressing red-shifted nuclear **a** mCitrine, **b** stagRFP, and **c** mCherry, and short-lifetime nuclear **d** slmGFP and **e** BruSLEE with 740 nm excitation. Detection in the NADH channel range 410-490 nm. Arrows indicate the transfected cells with nuclear FPs. Color is based on the phasor plot metabolic trajectories. Phasor plots associated with each FP are shown with metabolic arcs represented with a rainbow LUT to represent percent bound NADH, where blue is 0% and red is 100%. The left phasor represents the NADH channel that includes all pixels from the whole image. The right phasor plots the signal from the segmented nuclei of the transfected cells, indicated by the white arrows. Scale bars are 10 µm. Fluorescence intensity measurements from cytoplasm (orange line), untransfected control nuclei (blue line), and transfected nuclei (green line) of U2OS cells across excitation wavelengths from 680-780 nm with 10nm intervals in between. All measurements were normalized to excitation laser power (see Materials and Methods). Data points represent mean fluorescence intensities, and shaded regions indicate standard error of the mean. **a** BrUSLEE, **b** stagRFP, **c** mCherry, **d** slmGFP, and **e** mCitrine. Each data point represents the average of measurements from at least 5 nuclei per condition across multiple biological replicates.

In our hands, shifting the excitation wavelength of the FP did not allow for multiplexing with NADH. Therefore, we tested changing another fluorescence property, the fluorescence lifetime. All thus far tested green FPs have lifetimes on the order of 2-3 ns. We tested short lifetime monomeric GFP (slmGFP)^35^ and Bright Ultimately Short Lifetime Enhanced Emitter (BrUSLEE)^41^ with fluorescence lifetimes of 1.8 ns and 0.8 ns, respectively, in our TPE cross-talk assay (OPE spectra shown in **Supp. Fig. 7)**. At 740 nm, both FPs show weak fluorescence in the NADH channel, mapping close to the NADH cloud on the phasor (**Figure 4d,e**). The position of the signal on the phasor is more similar to the red-shifted FPs than to the other green FPs (shown in **Figure 2d-g**). TPE scans (**Figure 4a-c**) reveal that fluorescence levels are close to background for excitation <740 nm, but fluorescence in general (also at 940 nm TPE) is dim.

Recently, a set of monomeric, bright, and photostable fluorescent proteins based on StayGold (SG)^42^ have been developed. Their photobleaching behaviors are distinct from any other FP, implying novel photophysical properties. We chose mSG2^43^ and StayGoldE138D^44^ for assessment in our TPE assay, expressing and analyzing H2B fusions as previously. For both SG variants, we observe low TPE at 740 nm, negligible for StayGoldE138D, indicating that fluorescence intensities are comparable to those of untransfected nuclei (**Figure 5a and b**). However, phasor analysis of NADH and FP signals reveals overlap with NADH, similar to other green FPs.

**Figure 5:**
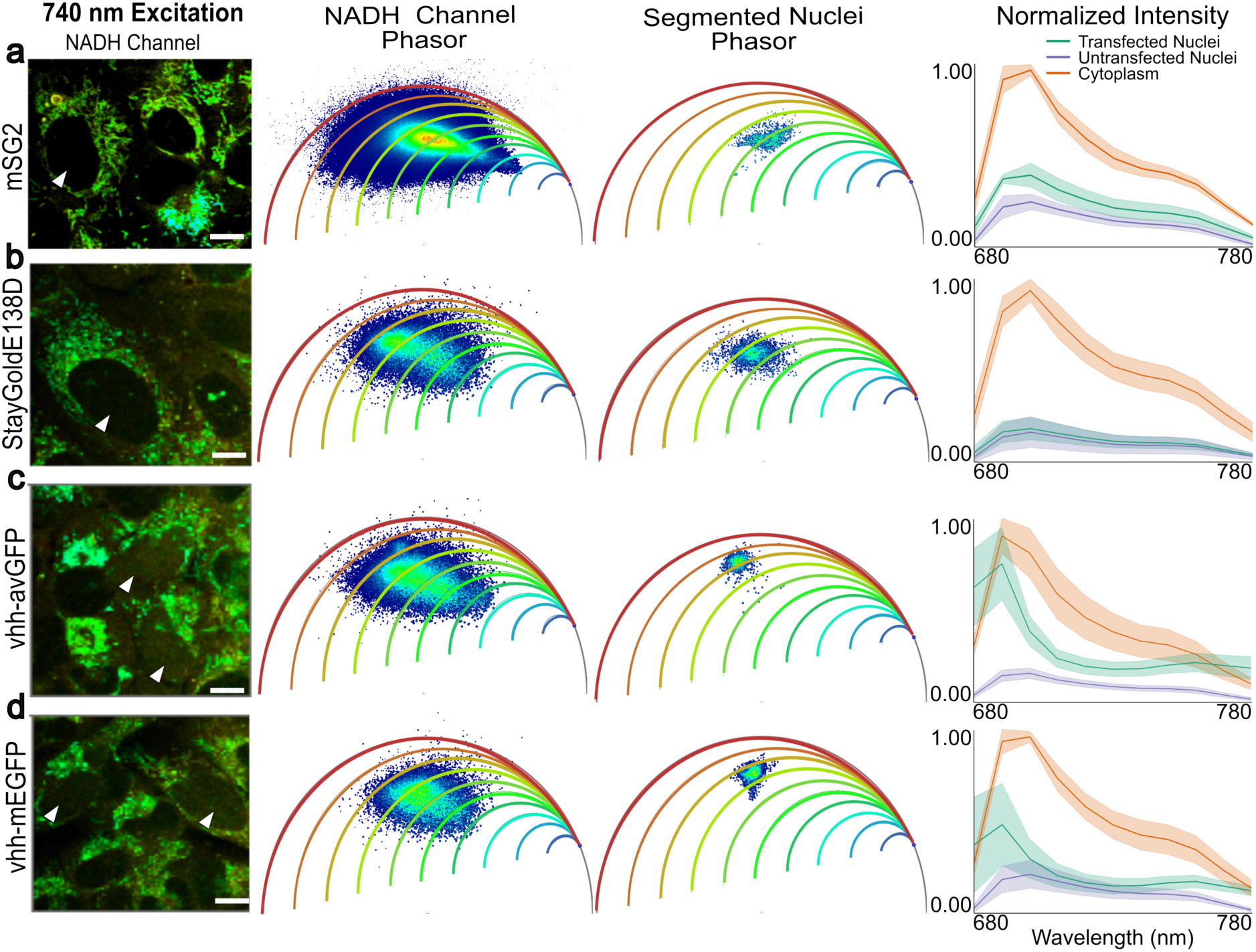
TPE cross-excitation assay using nuclear-localized FPs. Laser scanning TPE microscopy images of U2OS cells expressing engineered nuclear **a** mSG2, **b** StayGoldE138D, **c** vhh-avGFP, and **d** vhh-mEGFP with 740 nm excitation. Detection in the NADH channel range 410-490 nm. Arrows indicate the transfected cells with nuclear FPs. Color is based on the phasor plot metabolic trajectories below. Phasor plots associated with each FP are shown with metabolic arcs represented with a rainbow LUT to represent percent bound NADH, where blue is 0% and red is 100%. The left phasor represents the NADH channel that includes all pixels from the whole image. The right phasor plots the signal from the segmented nuclei of the transfected cells, indicated by the white arrows, determined using 940nm excitation images (not shown). Scale bars are 10 µm. Fluorescence intensity measurements from cytoplasm (orange line), untransfected control nuclei (blue line), and transfected nuclei (green line) of U2OS cells across excitation wavelengths from 680-780 nm with 10nm intervals in between. All measurements were normalized to excitation laser power (see Materials and Methods). Data points represent mean fluorescence intensities, and shaded regions indicate standard error of the mean. **a** mSG2, **b** StayGoldE138D, **c** vhh-avGFP, and **d** vhh-mEGFP. Each data point represents the average of measurements from at least 5 nuclei per condition across multiple biological replicates.

TPE at 740 nm, along with the resulting emission observed in the NADH channel and a shortened lifetime, suggests that the FP chromophore may decay from a specific state distinct from that typically exploited in standard imaging. The chromophores in the FPs can be protonated and deprotonated, which can have large effects not only on their absorption but also on their excitation spectra^45^. Changes in pH can be used to push the chromophore into one or the other state. Alternatively, certain nanobodies have been shown to enhance (E)GFP fluorescence by promoting the deprotonated state^33,46^. Building on the notion that a specific fluorescence pathway caused by a distinct chromophore state could cause the short lifetime fluorescence of the FPs observed in the NADH channel, we co-expressed the enhancer anti-GFP nanobody (vhh) with avGFP-H2B and EGFP-H2B. To ensure equimolar expression, we used the T2A multicistronic system. **Figure 5c-d** show the TPE Phasor-FLIM images, phasor plots, and excitation scans. TPE at 740 nm results in above background fluorescence for avGFP with a phasor position similar to avGFP without vhh (**Fig. 2f**). For vhh-mEGFP, TPE at 740 nm causes minimal fluorescence from the transfected nuclei compared with background from untransfected cells. Furthermore, in the presence of the vhh, a distinct, longer lifetime component becomes apparent on the phasor more similar to the phasor position for 940 nm-excited green FPs (**Fig. 2d**). However, some overlap with the metabolic NADH signal remains.

Metabolic signatures can be quite diverse and map all across the phasor plot space. Imaging untransfected cells reveals considerable heterogeneity, even within a single image (**Figure 6a,b**). To map out the expected expansion of the phasor, we used different drug and metabolic treatments to determine the biological minima and maxima for metabolic trajectory analysis in U2OS cells. Control cells, cells at low glucose (LG), high glucose (HG), and treated with Rot/AA, an ATP production inhibitor shutting down OXPHOS through complex I and III respectively, are shown in **Figure 6c**. During starvation with LG, cells shift towards OXPHOS. Upon HG stimulation, cells expectedly shift towards glycolysis. With HG + Rot/AA, two separate phasor populations emerge, one corresponding to the expected HG population and the other corresponding to the Rot/AA effect. As Rot/AA is known to inhibit OXPHOS, the cellular metabolism shifts towards glycolysis. Finally, we overlay that with the mean position of the segmented nuclei from transfected cells (**Figure 6d**). This summary analysis reveals that all tested FPs show a large degree of overlap with the NADH signature.

**Figure 6:**
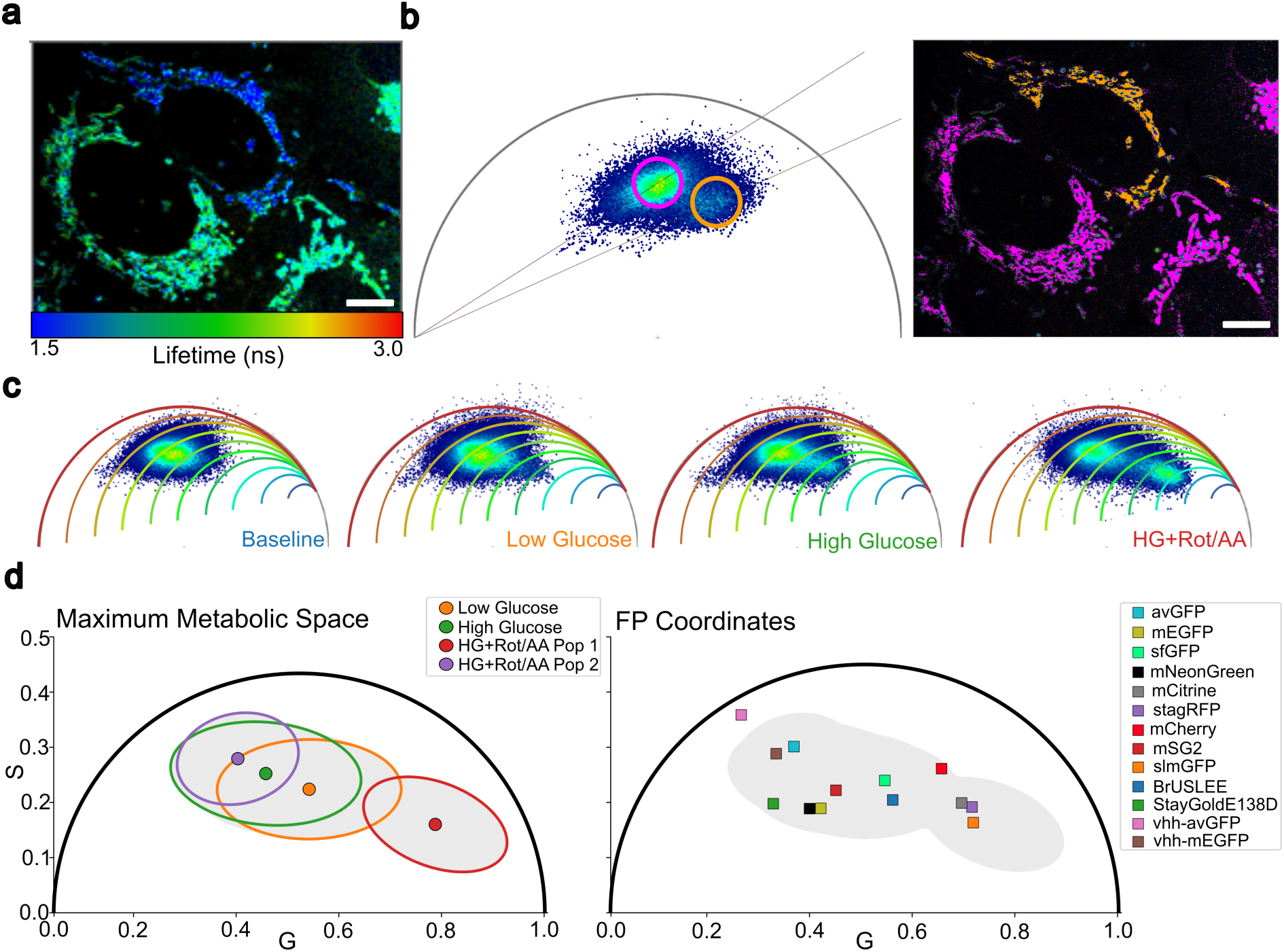
Maximum metabolic phasor and summary of the FP cross-excitation. **a** Laser scanning TPE microscopy image of untransfected U2OS with 740 nm excitation. Detection in the NADH channel range 410-490 nm. Arrows indicate regions of heterogeneous metabolic signal. Color is based on the lifetime (ns) with rainbow LUT, where blue is the shorter lifetime (1.5ns) and red is the longer lifetime (3.0ns). Scale bar 10µm. **b** Same image from **a** with color based on phasor plot below, where blue corresponds to the pixels within the blue circle and red to the red circle on the phasor plot to segregate heterogeneous cellular populations with different phasor metabolic signals. **c** Phasors from untransfected U2Os cells from Laser scanning TPE microscopy image with 740nm excitation and detection in NADH channel range of 410-490nm. Phasors are shown with metabolic arcs represented with a rainbow LUT to represent percent bound NADH, where blue is 0% and red is 100%. The phasor (from left to right) represents cells starved in low glucose (LG), cells stimulated with high glucose (HG), and the cells stimulated with high glucose and introduced to an oxidative phosphorylation inhibitor (HG+Rot/AA). **d** Metabolic trajectory boundary indicated through the black line determined by the pixels within two standard deviations from the respective mean G and S coordinates of each condition in **c**. LG is blue, HG is red, and HG+Rot/AA is green. Black line represents the pixels within a standard deviation of 2 from the respective means. **e** Mean G and S coordinate values of FPs plotted within metabolic trajectory boundary from **d**.

## Discussion

Metabolic imaging, exploiting TPE of NADH and the change in its fluorescence lifetime when bound to metabolic enzymes, is emerging as a powerful tool for studying metabolism in cells and tissues^14,18^. However, the exact acquisition conditions, for example, NADH TPE wavelength or detection, vary widely across the literature (**Table 3**). In this study, we try to provide a starting point for optimization, however, exact conditions will depend on the specifics of the biological sample being studied. Our primary focus is on the potential for multiplexing NADH metabolic imaging with other fluorophores, specifically fluorescent proteins.

Interpreting metabolic imaging data can be challenging and depends on a variety of controls, such as maximum OXPHOS and maximum glycolysis conditions measured for individual samples and studies. Multiplexing NADH FLIM with another label for specific cell types within a tissue or sub-cellular structures could aid in data interpretation and enable functional studies. Various FPs have been used previously in conjunction with NADH autofluorescence imaging. However, concerns have been raised that GFP and EGFP can be excited at 740 nm, causing a fluorescence signal indistinguishable from that of NADH in metabolic imaging^26,28,47^. These studies have not employed the powerful phasor approach to fluorescence, which has now become a standard tool to visualize metabolic imaging data. Our study combines the use of phasor FLIM and spatial separation of cytoplasmic NADH fluorescence from nuclear-localized FPs. While some NADH fluorescence is expected from the nucleus, it is significantly lower than that in the cytoplasm and predominantly comprises free NADH^48,49^. However, we need to acknowledge that the FP signal is likely being mixed with a low level of NADH autofluorescence, and thus in cells, we are quantifying fluorescence changes to this background.

Our experiments using recombinant FPs (**Supp Fig. S3, S4, and S6**) and cytoplasmic sfGFP and mNG (**Figure 1** and **Supp Figure 2**) confirm the reports of blue-shifted fluorescence under 740 nm TPE^26^. To quantitatively assess the effect and fully harness the power of the phasor transform, we move the FP signal to the nucleus using H2B fusions. The nuclear physicochemical environment differs from that of the cytoplasm; yet we observe similar fluorescence properties in cyt-mNG/sfGFP compared with H2B-mNG/sfGFP. Strikingly, under NADH imaging conditions (740 nm excitation and 410-490 nm detection), we record increased intensity and shortened fluorescence lifetime compared with untransfected nuclei within the same sample. The excitation and fluorescence are weak compared to 940 nm TPE, however, since the signal from NADH at 740 nm is also weak, it may still pose an obstacle to effective multiplexing.

We observe the unusual excitation at 740 nm and fluorescence within the NADH detection window for several widely used FPs across the visible spectrum. Varying the excitation wavelengths changes the effect slightly, but we could not identify a condition where it is abolished completely. Fluorescence at 740 nm TPE from red-shifted FPs such as mCherry and stagRFP was unexpected and suggests that the observed fluorescence does not originate from cross-talk of the typical fluorescence peak into the NADH channel. Instead, a different state is excited or induced, similar to what has been suggested for dsRed^28,50^. At this point, we cannot exclude that the 740 nm laser induces a photo-conversion or color change on the FPs. However, we did not notice an increase in the blue-shifted signal over time. Among the 11 tested FPs, the most promising candidates for multiplexing are mNG, mSG2 and StayGoldE138D. These FPs show negligible excitation at 740 nm and have the benefit of high photostability under OPE^43,44,51^.

The analysis of the TPE spectra in live cells shows that slightly varying the excitation wavelength does not allow for mitigation of the cross-excitation. A limitation of our assay is that we have no control over the expression levels of the FP-H2B proteins as we were employing transient transfections. Variability in expression explains the large standard deviation in the excitation scans, which are based on fluorescence intensity. Using stable expressions could mitigate that but we would not expect a change to our observations and conclusions as fluorescence lifetime is robust against changes in concentration^52^.

What causes the unusual fluorescence properties of the FPs under 740 nm remains a matter of speculation. We are getting some hints from the co-expression of mEGFP and avGFP with the enhancer nanobody (vhh). The vhh is known to shift the FP’s chromophore to the deprotonated state, thus forcing a specific fluorescence decay route. In our study, this seems to reduce the emission in the NADH channel and shifts it to the FP channel. We co-expressed the nanobody and FP using a T2A system. A tandem version could potentially be more efficient but bears the risk of oligomerization.

Our findings implicate challenges to multiplex NADH with fluorescent protein reporters. We aim to raise awareness that cross-excitation represents a widespread phenomenon, influencing fluorescent proteins across the visible spectrum rather than merely green variants.. We strongly recommend the spatial segregation of FP signals from endogenous NADH; for instance, directing FPs to the membrane or the nucleus (as shown here with H2B fusions effectively isolates the two signals). Careful analysis and controls are paramount: evaluating untransfected and transfected cells within the same field of view allows for the identification and, possibly, the subtraction of cross-talk contributions. While stable, uniform FP expression may permit the use of constant background corrections to yield reliable metabolic data, variable expression, prevalent in transient transfections or rapidly developing tissue, introduces cell-to-cell intensity fluctuations that impede measurements of metabolic signatures, potentially inducing systematic errors. Consequently, multiplexed strategies in these contexts require extreme caution. Among the FPs evaluated in this work, StayGoldE138D and mNeonGreen are identified as the most compatible candidates; both demonstrate negligible excitation in the NADH channel under 740 nm TPE and are thus preferable to standard alternatives such as mCherry.

Overall, we describe the observation that multiplexing metabolic imaging of NADH at 740 nm TPE with fluorescent proteins is challenging. This is a cautionary tale that provides some insights into the underlying mechanisms and issues. We provide a path towards a solution and remain carefully optimistic that with the right controls and spatial separation of the signals, multiplexing of NADH and FPs will be possible.

## Acknowledgments

F.S. acknowledges support from the long-term postdoctoral fellowships from EMBO (EMBO ALTF 849-2020) and HFSP (LT000404/2021-L) as well as from the Leverhulme Trust Leverhulme Trust, UK (LIP-2021-017). O.A. acknowledges support from the USC NIH T32 PhD Fellowship in Developmental Biology, Stem Cells, and Regeneration (5T32HD060549). J.C. acknowledges support from the ARCS foundation and CHLA TSRI Core Pilot Grant. L.C. acknowledges support and guidance from Stacey D. Finley. The authors thank Tristan Nicholas (University of Warwick) for critical proofreading of the manuscript. Jason Junge and Arkadi Shwartz at the Translational Imaging Center (USC) for technical support and maintenance of the Leica SP8.

## Author Contributions

Conceptualization, F.S.; Methodology, F.S., J.C., and O.A.; Software, F.S. and J.C.; Formal Analysis, J.C., O.A., and L.C.; Investigation, F.S., J.C., O.A., Y.Z.P., and L.C.; Resources, O.A., Y.Z.P., and L.C.; Data Curation, T.S., C.D., and A.V.; Visualization, J.C., O.A., and L.C.; Writing – Original Draft, F.S.; Writing – Review & Editing, F.S., J.C., O.A., L.C., K.W., S.E.F., S.K.G., and S.K.; Supervision, F.S., K.W., S.E.F., S.K.G., and S.K.; Funding Acquisition, K.W., S.E.F., S.K.G., and S.K.

## Code and Data Availability

All code used to analyse data is available at https://github.com/janiellecuala/Multiplexing-NADH-and-Fluorescent-Proteins-in-Metabolic-Imaging.

All raw data can be requested from the corresponding author upon reasonable request.

## Notes

### Competing Interest Statement

The authors have declared no competing interest.

