## Supporting Figures for "Bright Probes, Blurred Metabolism: Navigating Fluorescent Protein Cross-Excitation in NADH FLIM"

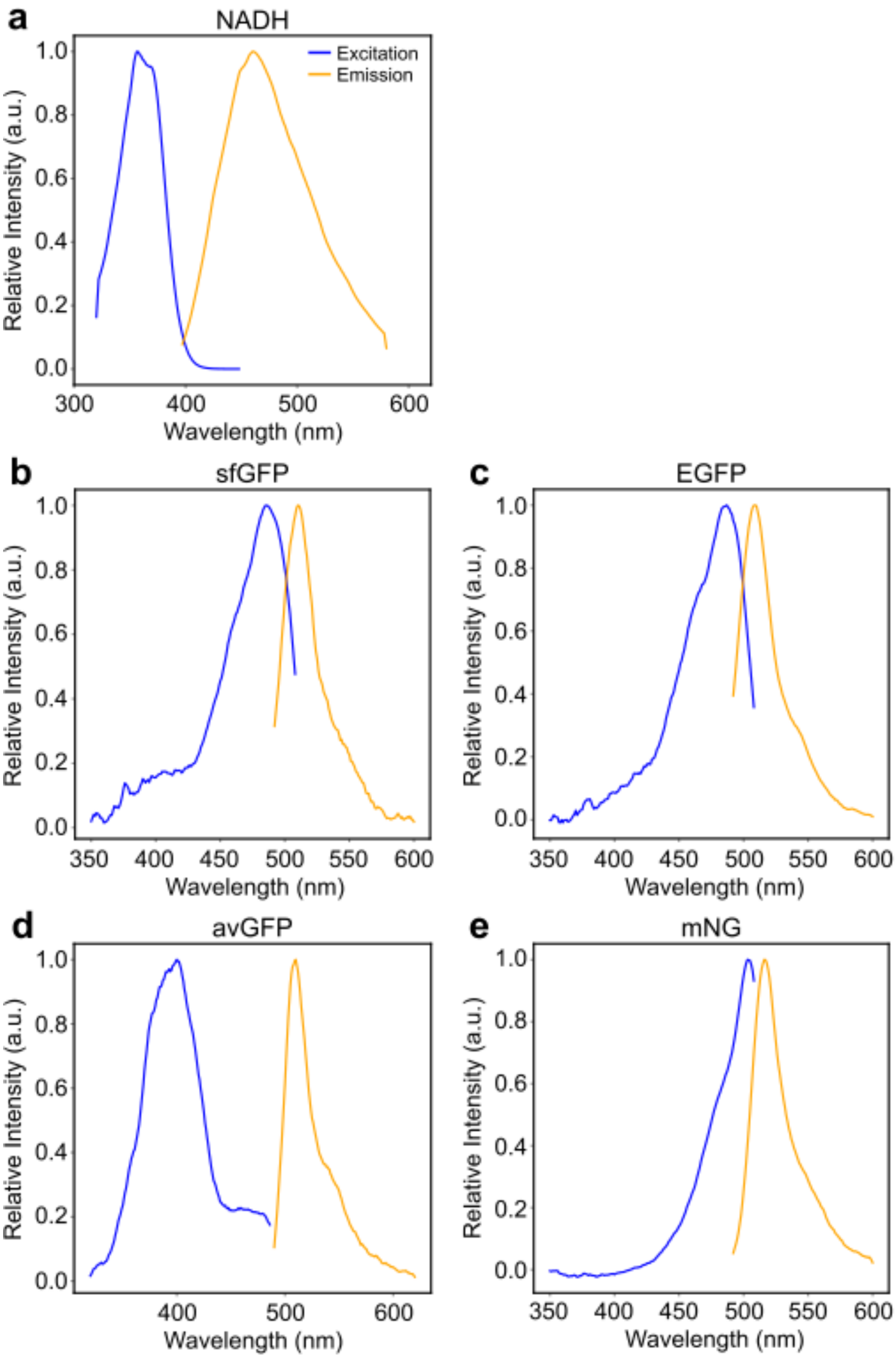


**Supplementary Figure S1:** One photon excitation and emission spectra of NADH in solution and green fluorescent proteins expressed in U2OS cells as H2B fusions. Blue curves are the excitation spectra, and the yellow curves are the emission spectra.

**a** NADH in water

**b** sfGFP in cells

**c** mEGFP
**d** avGFP
**e** mNG
Spectra were acquired in glass bottom 96 well plates on a fluorescence plate reader. Lines represent averages of multiple repeats.

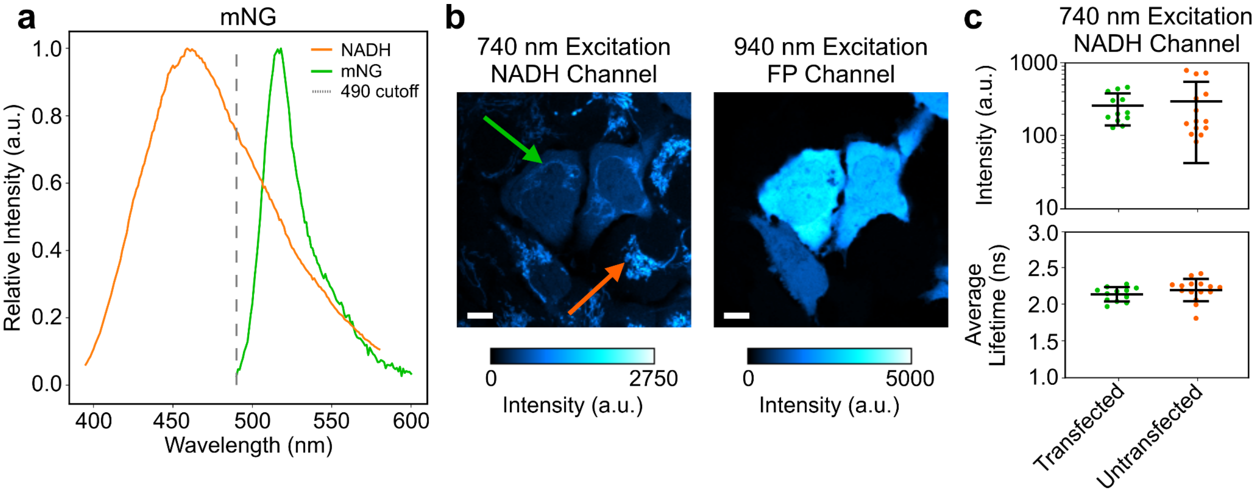


**Supplementary Figure S2:** Cross-talk of NADH and mNeonGreen (mNG).

**a** Spectral overlap of NADH (in water) and mNG-H2B expressed in HEK cells

**b** Laser scanning TPE microscopy images of cells expressing cytoplasmic mNG. Left: Excitation with 740 nm. Right: Excitation with 940 nm. Arrows indicate mitochondria in transfected (green) and untransfected (orange) cells. Scale bars are 10 µm.

**c** Fluorescence intensity and lifetime of segmented cytoplasm in transfected (green) and untransfected (orange) cells. Every dot represents one cell. Horizontal bar is mean value and vertical bars standard deviation.

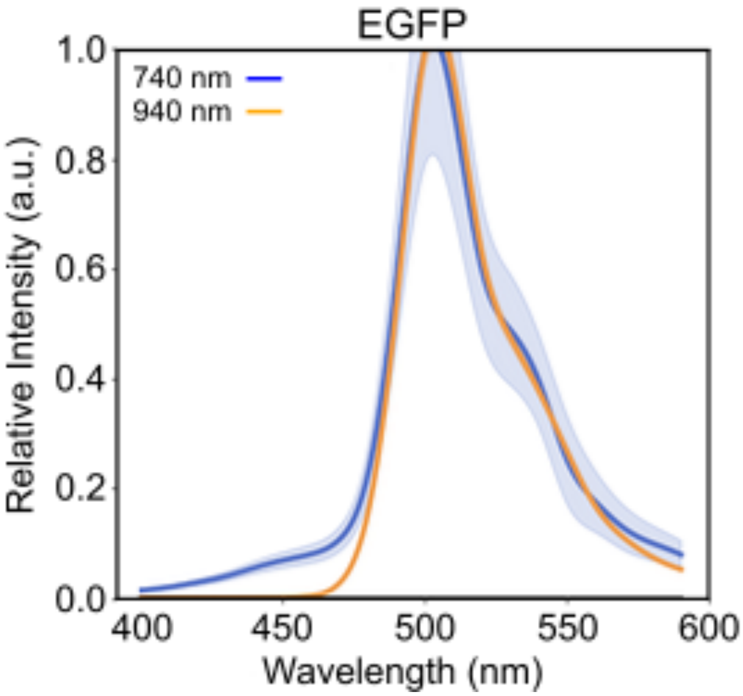


**Supplementary Figure S3:** Two-photon emission spectra of recombinant EGFP in L-15. 150 nM solution of EGFP in L-15 was excited using 740 nm (blue) and 940 nm (orange). Values are averaged across an image.

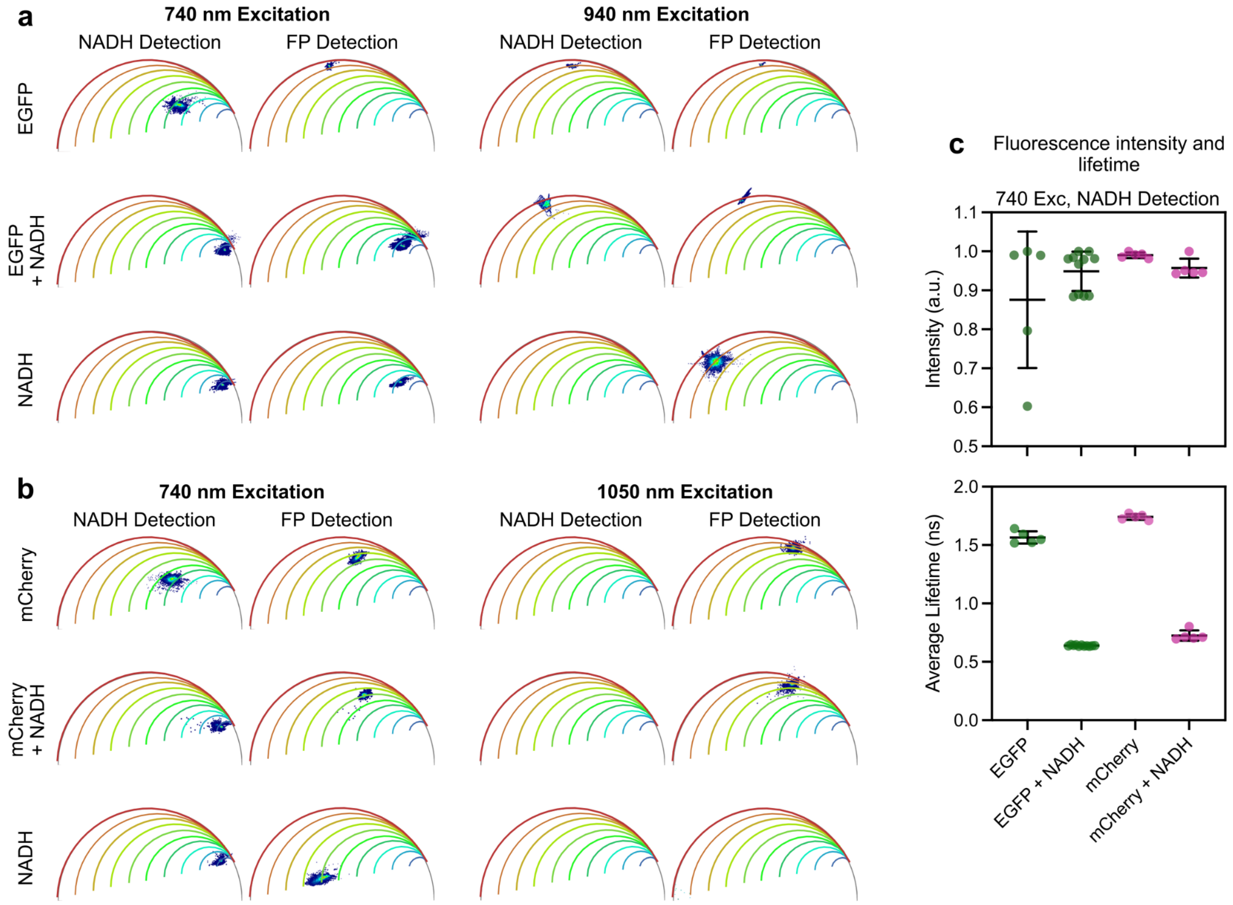


**Supplementary Figure S4:** Excitation of purified recombinant EGFP and mCherry in the presence and absence of NADH in L-15. EGFP is imaged at a concentration of 150 nM, mCherry is imaged at 250 nM, and NADH is imaged at 200 μM.

**a** Phasors for recombinant EGFP and NADH fluorescence lifetime. Left: Excitation with 740 nm and NADH detection (410-490 nm, left) and FP detection (500-600 nm, right). Right: Excitation with 940 nm and NADH detection (left) and FP detection (right).

**b** Phasors for recombinant mCherry and NADH fluorescence lifetime. Left: Excitation with 740 nm and NADH detection (410-490 nm, left) and FP detection (600-700 nm, right). Right: Excitation with 1050 nm and NADH detection (left) and FP detection (right).

**c** Fluorescence lifetime and intensity of recombinant FPs with 740 nm excitation and NADH detection. Horizontal bars indicate mean value and vertical bars indicate standard deviation.

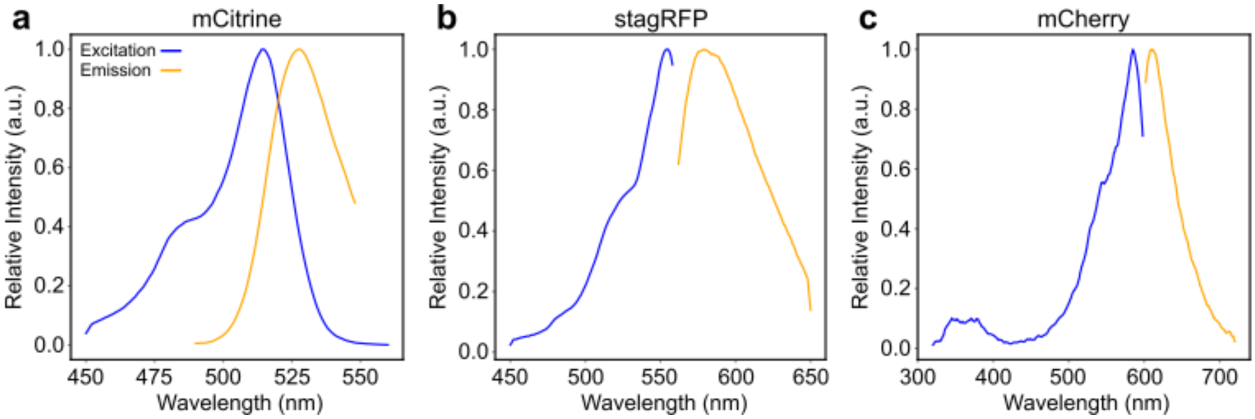


**Supplementary Figure S5: One photon excitation spectra of yellow and red-shifted FPs**
Fluorescent proteins expressed in U2OS cells as H2B fusions. Blue curves are the excitation spectra, and the yellow curves are the emission spectra.

**a** mCitrine

**b** stagRFP

**c** mCherry

Spectra were acquired in glass bottom 96 well plates on a fluorescence plate reader. Curves represent averages of multiple repeats.

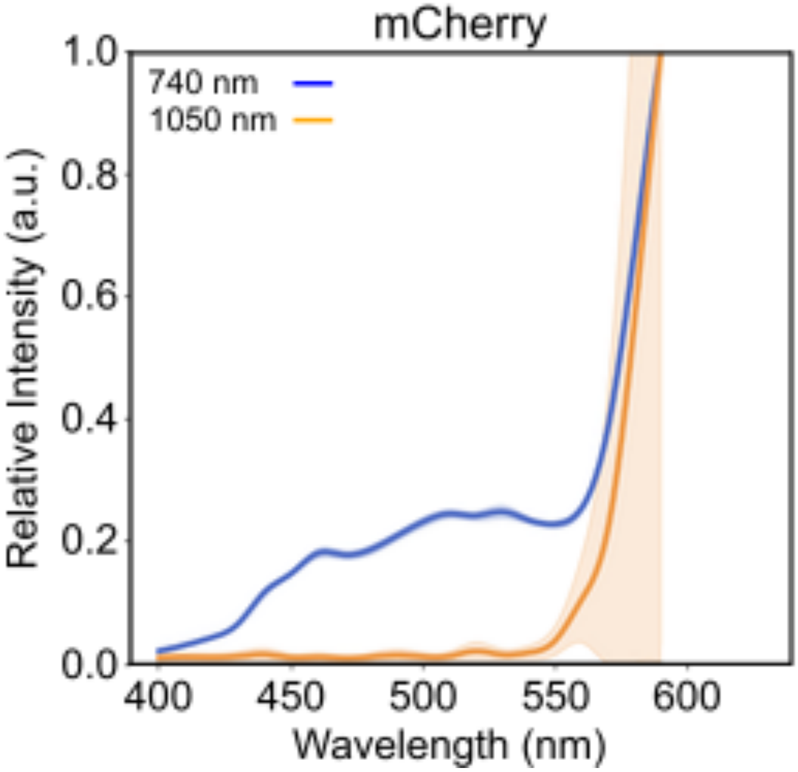


**Supplementary Figure S6:** Two-photon emission spectra of mCherry at 740 nm and 1050 250 nM solution of mCherry in L-15 was excited using 740 nm (blue) and 1050 nm (orange). Values are averaged across an image and normalized to the maximum value.

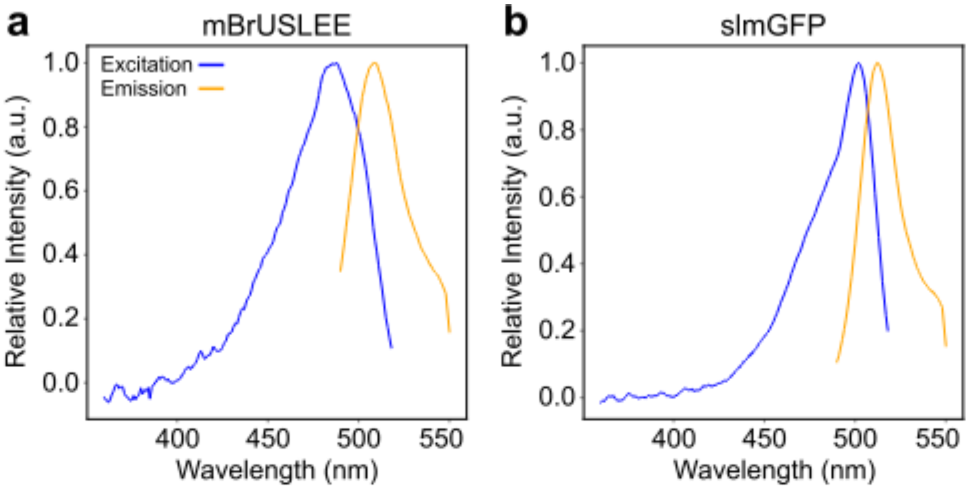


**Supplementary Figure S7: One-photon excitation spectra of short fluorescence lifetime FPs**

Fluorescent proteins expressed in U2OS cells as H2B fusions. Blue line is the excitation spectrum, and the yellow line is the emission spectrum.

**a** mBrUSLEE

**b** slmGFP

Spectra were acquired in glass bottom 96 well plates on a fluorescence plate reader. Lines represent averages of multiple repeats.
